# Transcriptome profiling of maize transcription factor mutants to probe gene regulatory network predictions

**DOI:** 10.1101/2024.07.30.605884

**Authors:** Erika L. Ellison, Peng Zhou, Yi-Hsuan Chu, Peter Hermanson, Lina Gomez-Cano, Zachary A. Myers, Ankita Abnave, John Gray, Candice N. Hirsch, Erich Grotewold, Nathan M. Springer

## Abstract

Transcription factors (TFs) play important roles in regulation of gene expression and phenotype. A variety of approaches have been utilized to develop gene-regulatory networks (GRNs) to predict the regulatory targets for each TF, such as yeast-one-hybrid (Y1H) screens and gene co-expression network (GCN) analysis. Here we identified potential TF targets and used a reverse genetics approach to test the predictions of several GRNs in maize. Loss-of-function mutant alleles were isolated for 22 maize TFs. These mutants did not exhibit obvious morphological phenotypes. However, transcriptomic profiling identified differentially expressed genes in each of the mutant genotypes, and targeted metabolic profiling indicated variable phenolic accumulation in some mutants. An analysis of expression levels for predicted target genes based on Y1H screens identified a small subset of predicted targets that exhibit altered expression levels. The analysis of predicted targets from GCN-based methods found significant enrichments for prediction sets of some TFs, but most predicted targets did not exhibit altered expression. This could result from false-positive GCN predictions, a TF with a secondary regulatory role resulting in minor effects on gene regulation, or redundant gene regulation by other TFs. Collectively, these findings suggest that loss-of-function for single uncharacterized TFs might have limited phenotypic impacts but can reveal subsets of GRN predicted targets with altered expression.

## Introduction

Transcription factors (TFs) transcriptionally regulate gene expression by recognizing and binding to DNA in a sequence-specific fashion. In eukaryotic genomes ∼5-10% of the genes encode TFs that regulate transcription of all genes (Riechmann 2002; Sperling 2007; Brkljacic and Grotewold 2017; Lambert *et al*. 2018). Gene regulatory networks (GRNs) represent the interactions between TFs and target genes that regulate spatial and temporal expression of genes (Macneil and Walhout 2011; Mejia-Guerra *et al*. 2012; Badia-I-Mompel *et al*. 2023). The data to match the discrete number of TFs to the larger number of target genes they regulate in GRNs remains limited. However, it is important to identify and understand how GRNs regulate endogenous metabolic pathways as this may provide key insights for modulating whole pathways or branch points of pathways (Farré *et al*. 2014). Likewise, GRN inference can be used to select for existing variants or introduce novel TF alleles as a potential strategy to generate novel phenotypes for crop improvement (Springer *et al*. 2019).

Several methods have been used to predict TF-target gene interactions to generate GRNs in maize. These methods can include gene-centered approaches; a gene is known but regulators of the gene are not, or TF-centered approaches; the TF is known but the target genes it regulates are not (Yang *et al*. 2016). Gene-centered approach methods can identify interactions where TFs directly bind to promoters or *cis*-regulatory elements (CREs) of a particular gene. Yeast-one-hybrid (Y1H) is a gene-centered approach that involves screening for interactions between the DNA sequence of interest (DNA bait) and a TF (protein prey) by activation of a reporter gene in yeast (Arda and Walhout 2010; Yang *et al*. 2016). Studies in *Arabidopsis* (Brady *et al*. 2011; Taylor-Teeples *et al*. 2015; Ikeuchi *et al*. 2018) and maize (Yang *et al*. 2017; Abnave *et al*. 2024) have successfully used Y1H to predict TF-target gene interactions. However, there are numerous limitations of the Y1H approach, including interactions are tested in yeast outside of genome tissue-specific or chromatin landscapes, cloned promoter sequences (DNA bait) are usually short (∼1 kb) and may not capture the TF binding site, and interactions that require multiple TFs or post-translational modifications will be missed (Arda and Walhout 2010; Walhout 2011). TF-centered approaches, such as chromatin-immunoprecipitation coupled with DNA sequencing (ChIP-seq), can identify potential TF targets genome-wide. ChIP-seq is a technique that uses a TF-specific antibody to selectively recover bound DNA from cross-linked DNA-protein complexes (Kuo and Allis 1999; Johnson *et al*. 2007). In maize, direct targets for several TFs have been identified by ChIP-seq (Morohashi *et al*. 2012; Bolduc *et al*. 2012; Eveland *et al*. 2014; Pautler *et al*. 2015; Li *et al*. 2015; Tu *et al*. 2020; Hartwig *et al*. 2023). Unlike Y1H, ChIP-seq captures *in vivo* TF-target interactions within accessible chromatin regions and identifies target sequences that are both directly and indirectly bound (if the TF of interest forms a complex with another TF that binds the target directly). ChIP-seq is limited by the availability of antibodies and loss-of-function alleles to test antibody specificity. Another approach to GRN inference is to build gene co-expression networks (GCNs) by statistically measuring the relationship between TF and target gene expression profiles. If the expression pattern of the TF and target gene are similar, these genes are considered co-expressed and may have shared regulation (Eisen *et al*. 1998). GCNs require large sets of quantitative data, usually RNA-seq, to capture gene expression and the statistical methods used for correlation can have a significant impact on the results (Huang *et al*. 2017). Each approach to predict GRNs could identify portions of true GRNs, but comparisons between these predictions reveal only limited overlap, suggesting many false positives and false negatives from each approach.

One method to test GRN predictions is to isolate loss-of-function mutants in TFs and test expression of predicted targets (Scherens and Goffeau 2004; Thompson *et al*. 2015). In maize, there are limited methods for moderate scale reverse genetics studies to assess if absence of the TF results in target genes with altered expression. Current maize mutant libraries only provide functional knockouts for a subset of genes in the genome (Lu *et al*. 2018). Further, the ability to test GRN predictions with TF loss-of-function alleles will vary based on the GRN prediction method. For example, the sample size of putative TF regulators identified by GCN-based methods is usually much larger than those identified by Y1H screens. The number of Y1H predicted TF-target interactions is limited by the size and number of promoter regions that are cloned and tested, usually from a single putative pathway or functional type. GCN-based methods allow for construction of much larger networks and the likelihood of isolating loss-of-function alleles for some of these predicted TF regulators increases by the size of the network alone. Due to the recent whole genome duplication in maize (Gaut *et al*. 2000), testing functional impacts of GRN predictions *in vivo* with TF knockouts may be limited by genetic redundancy, which requires loss-of-function mutants for multiple related TF genes.

A set of putative loss-of-function alleles were recovered for 22 maize TFs with predicted GRN targets based on either Y1H and/or GCNs. We did not observe major phenotypic differences in these mutants under normal growth conditions. However, transcriptome profiling revealed variable numbers of differentially expressed (DE) genes in transposon-insertion mutants relative to the corresponding inbred line plants. The analysis of transcript abundance for GRN targets predicted by Y1H or GCN methods does reveal examples of predicted targets with altered expression. In some cases, the predicted targets are significantly over-represented in the DE genes. However, the majority (> 75%) of predicted targets did not exhibit altered expression. Gene ontology (GO) enrichment analyses identified functional groups of genes with altered expression in some of these mutants but did not point to clear biological functions for these TFs. These findings suggest limited perturbation of GRNs in these single gene TF knockouts and could reflect high degrees of functional redundancy in gene regulation.

## Results

To test the functional relevance of GRN predictions in maize we obtained stocks containing putative loss-of-function alleles for a series of TFs. Two primary sources of GRN predictions were utilized to select TFs for testing. The first source of GRN predictions was from a Y1H screen that identified putative TF regulators of maize phenolic biosynthesis (Yang *et al*. 2017)). This Y1H screen identified 45 TFs that exhibit interactions with at least 4 of the 54 phenolic biosynthesis gene promoters tested (Burdo *et al*. 2014; Yang *et al*. 2017). The second source of GRN predictions was a meta-analysis of TF-target gene co-expression from 45 GCNs (Zhou *et al*. 2020b). To test the GRN predictions generated from these 45 GCNs, we identified 64 TFs that had ≥ 400 predicted targets and at least one coding sequence insertion indexed in the UniformMu mutant collection (McCarty *et al*. 2005). GRNs constructed from both sources, Y1H and GCNs, identified predictions for genes annotated in the B73v4 genome (Schnable *et al*. 2009; Jiao *et al*. 2017). Before moving forward with network perturbation analyses, all TFs selected for testing (45 from Y1H and 64 from GCNs) were confirmed to be single copy syntenic orthologs in the B73v4 and W22 (considered the wild type in this study) genomes (Monnahan *et al*. 2020).

### Isolation of TF mutant alleles from the UniformMu population

*Mutator* (*Mu*) transposon insertions located within the set of TFs predicted from the Y1H or GCNs were identified using the sequence-indexed UniformMu population created in a W22 inbred genetic background (McCarty *et al*. 2005). While most maize genes are associated with *Mu* insertions in this population, only 27.1% of annotated W22 genes have an insertion within the coding sequence (Springer *et al*. 2018). We identified all available insertions for the 45 TFs identified as candidates in the Y1H screen, including insertions in UTRs, introns, and proximal promoter regions (up to 1 kb upstream of the annotated TSS). Given the prior evidence that some insertions within UTRs, introns, or 5’ regions have minimal or no effect on the transcript produced by the allele (Kidwell and Lisch 1997; Dietrich *et al*. 2002; Liu *et al*. 2009), we focused primarily on coding region insertions for GCN predicted TFs. We initially obtained stocks representing 150 alleles (82 TFs) but some of these were subsequently eliminated from our study because we could not confirm the presence of the insertion or there was lack of evidence for loss-of-function. In total, we isolated 32 putative loss-of-function alleles for 22 TFs, including 12 alleles for 8 TFs selected from initial Y1H predictions (Yang *et al*. 2017) and 20 alleles for 14 TFs selected from GCN predictions (Figure 1, Table S1). After isolating the mutant alleles for the GCN selected TFs, another Y1H screen was performed using the promoters of the top 19 predominantly expressed phenolic genes from Gomez-Cano et al. 2020 (Gomez-Cano *et al*. 2020) as bait, 7 of which were not included in the initial Y1H screen with 54 phenolic promoters (Abnave *et al*. 2024). This new Y1H screen identified phenolic targets for 6 of the TFs with GCN predicted targets (Abnave *et al*. 2024). In total, 13 of the 22 TFs in this study were predicted to regulate phenolic biosynthesis genes by Y1H (Figure 1, Table S1).

**Figure 1.**
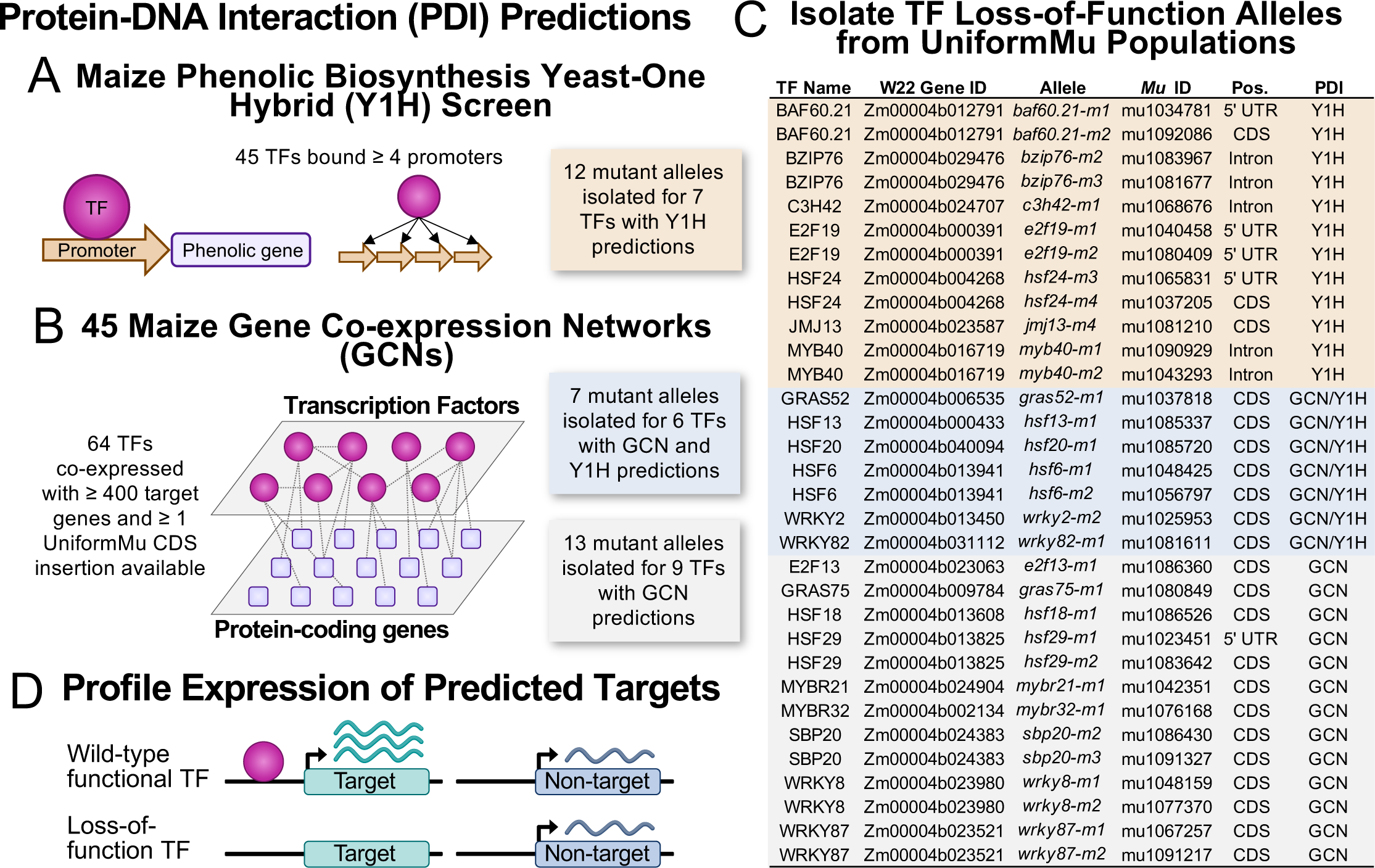
Experimental design to test transcription factor-target gene predictions in maize. Maize transcription factors (TFs) were predicted to be transcriptional regulators of a set of target genes based on **(A)** a maize phenolic biosynthesis Y1H screen (Yang *et al*. 2017) or **(B)** construction of 45 maize gene co-expression networks (GCNs) (Zhou *et al*. 2020). To test these TF-target gene predictions in vivo we focused on **(A)** 45 TFs that bound ≥ 4 phenolic biosynthesis promoters in the Y1H screen or **(B)** 64 TFs that were co-expressed with ≥ 400 target genes across the 45 GCNs and had an available UniformMu insertion in the TF gene coding sequence. **C)** The UniformMu population was used to isolate 12 mutant alleles for 7/45 Y1H predicted TFs and 20 mutant alleles for 15/64 GCN predicted TFs (Table S1). 7 of the 15 GCN predicted TFs with mutant alleles were also predicted to be phenolic regulators in an additional Y1H screen (Abnave *et al*. 2024). **D)** Transcriptome profiling was completed on wild-type plants with a functional TF and mutant plants to quantify expression of predicted target genes.

### Identifying TF loss-of-function alleles and DE genes by transcriptome profiling

Transcriptome profiling by RNA-seq was performed for each TF mutant allele to characterize genome-wide perturbations of expression. We sampled a single tissue for each TF mutant in which the TE gene exhibited moderate expression. Across the 22 TFs, a total of five different tissues were sampled (Figure S1, Table S1, Table S2). To confirm the potential functional impact of the *Mu* insertion on the gene product of the TF, we assessed the expression level and transcript structure for each mutant allele. The change in mRNA accumulation for each TF gene in the mutant allele harboring the *Mu* insertion relative to the wild type allele was estimated from RNA-seq reads of all biological replicates mapped to the W22 reference genome (which lacks the *Mu* insertion). This expression analysis revealed that 8 of the 32 mutant alleles had significantly reduced expression levels and 2 (*wrky8-m1* and *hsf18-m1*) exhibited significant increases in total transcript abundance (Figure 2, Table S1) (Ellison *et al*. 2023). However, the lack of a difference in mRNA accumulation level does not necessarily mean that a functional product is produced. To determine if the *Mu* insertion resulted in potential loss-of-function alleles, we generated a *de novo* transcript assembly and identified variation in transcript structure and sequence for each mutant. Our prior work found that most mutant alleles have either altered transcript structure or sequence that is predicted to result in the production of truncated or altered protein sequences (Ellison *et al*. 2023). In this study, we only retained mutant alleles that could not produce the full-length protein by selecting mutants with either transcripts 5’ of the *Mu* insertion with the initial AUG out-of-frame and/or transcripts 3’ of *Mu* that would only produce a protein less than half the length of the normal protein. Based on the assembled transcripts, it is unlikely that functional proteins are produced for most of these mutant alleles, although it is possible that partial fragments could be generated in some cases.

**Figure 2.**
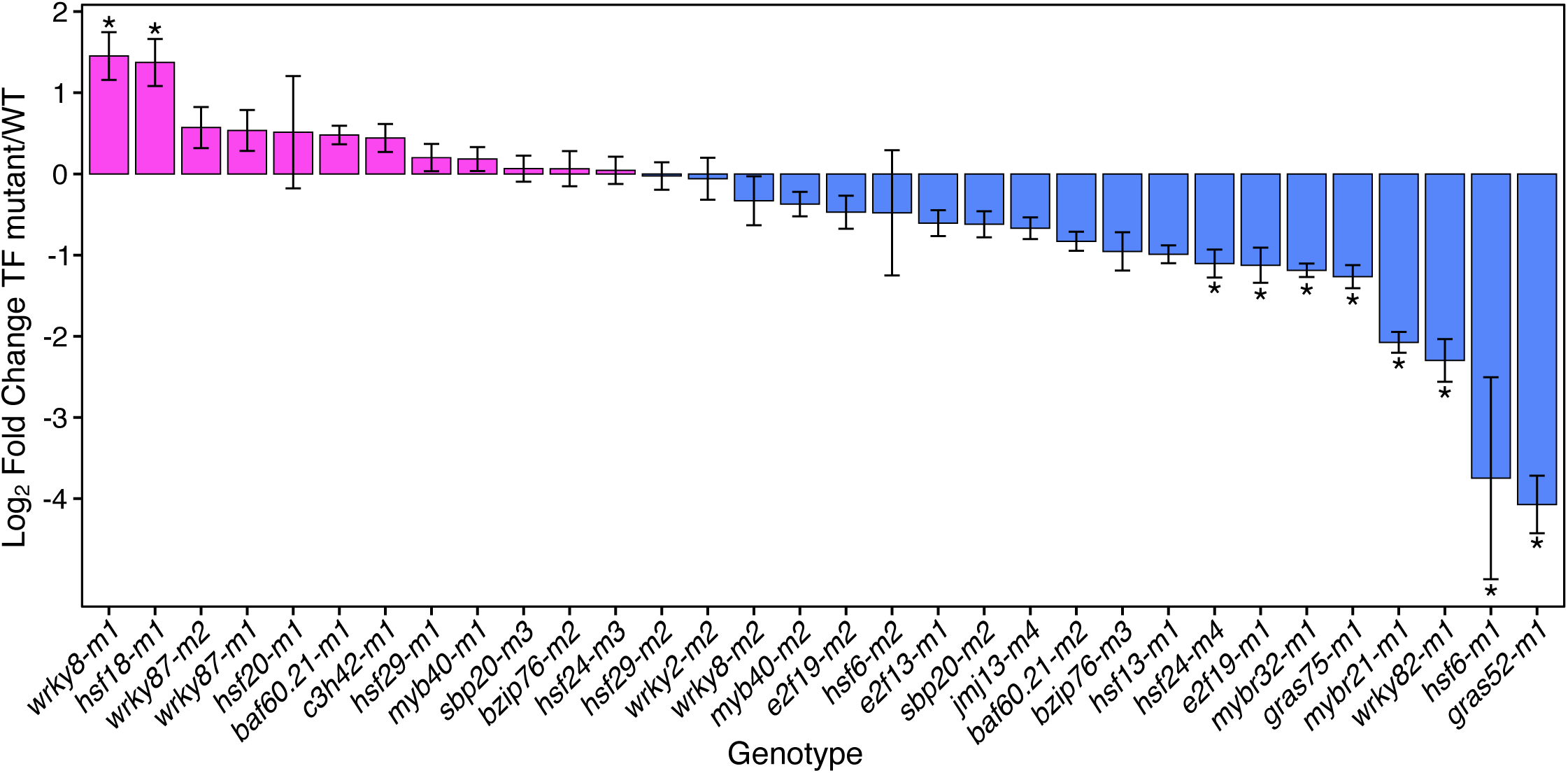
Changes in transcript abundance for TF mutant alleles. The fold change in expression of each TF gene in the mutant allele harboring the *Mu* insertion relative to wild type plants was estimated from RNA-seq reads of all biological replicates mapped to the W22 reference genome (log2 fold change TF mutant allele/WT). The standard error of the log2 fold change (log2fc) estimate is represented as error bars for each allele. Significant differential expression between the TF mutant and W22 control is indicated by an asterisk (*). Alleles are rank ordered based on the difference in expression for this plot with positive log2fc: pink and negative log2fc: blue.

The overall changes to the transcriptome were assessed through principal component analysis for each tissue (Figure S1). In general, the samples tended to cluster by mutant allele, but some samples were more like W22 wild type while others were more distinct (Figure S1). Genes that were DE were identified for each mutant allele relative to W22 replicates from the same tissue (Figure 3). The number of DE genes (DEGs) was quite variable with some mutants only exhibiting ∼100 and others having > 1,500 (Figure 3, Data File S1). For these putative TFs it is not known whether they function as primarily activators or repressors; therefore, both up- and down-regulated genes were identified for each mutant allele. Most of the mutants (26/32) have more down-regulated genes than up-regulated genes, which would be expected for TFs that have a primary role as transcriptional activators. We examined the 10 TFs that are represented by two independent mutant alleles and found significant overlap in both the up- and down-regulated DEGs for all 10 pairs of mutant alleles. While the overlap of DEGs between the two mutant alleles was highly significant, the proportion of the up- or down-regulated genes that are significant in both mutant alleles was highly variable (Figure S2). In several cases, > 50% of the DEGs are identified in both mutant alleles, but in other cases the overlap only accounted for 5-10% of the DEGs (Figure S2).

**Figure 3.**
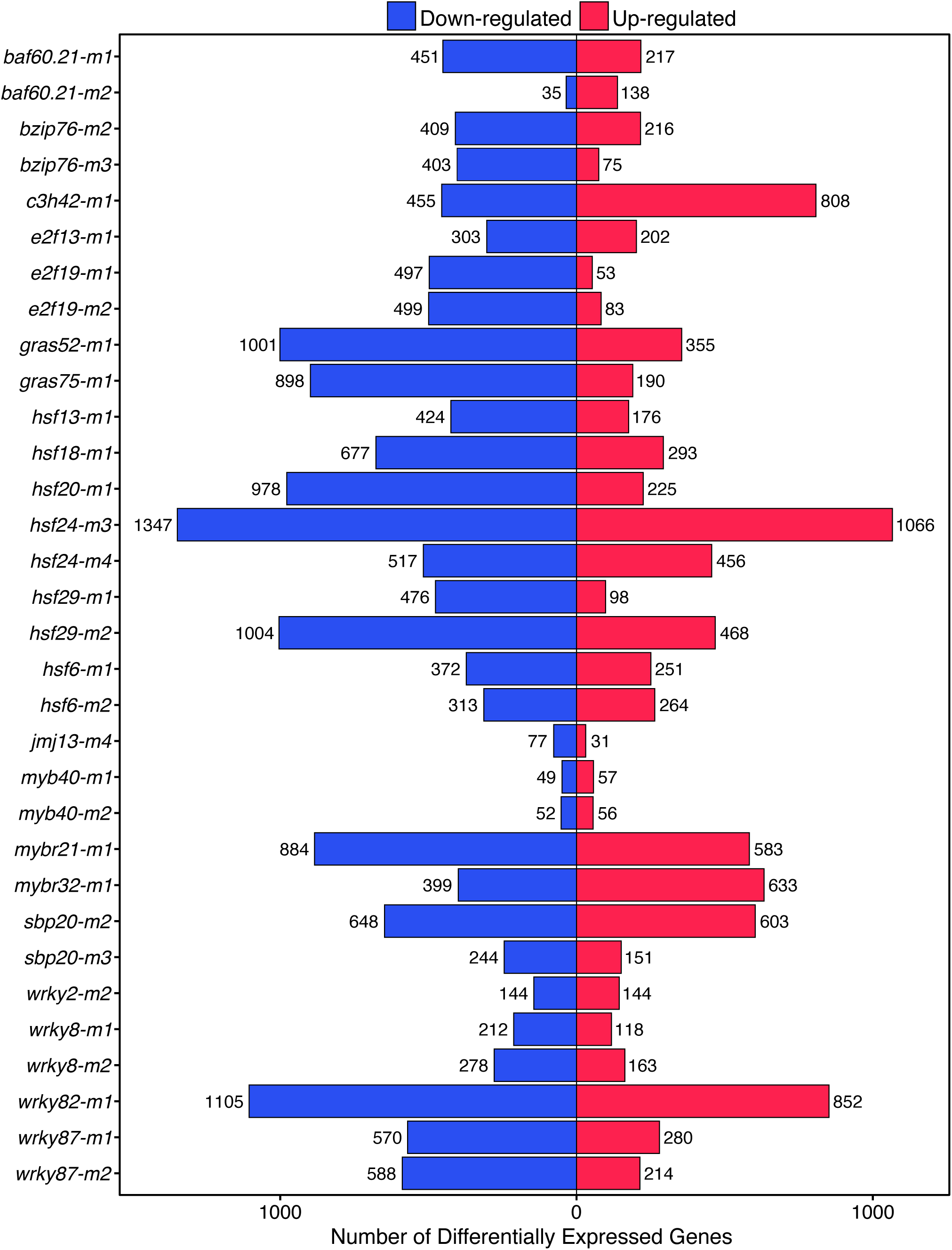
Identification of differentially expressed genes in transcription factor mutant genotypes. Genes with significant differences in expression were determined for each mutant based on comparison to W22 samples of the same tissue type using DEseq2 (FDR adjusted p < 0.05 and fold-change > 2). The number of significantly up- (red) or down-regulated (blue) differentially expressed genes (DEGs) in the TF mutant allele relative to the W22 control is shown.

### Morphological and metabolic phenotypic characterization of TF mutant lines

There were no obvious visible abnormalities observed in any of the TF mutant lines. To determine if the TF mutant lines exhibited any quantitative variation we measured two plant architecture traits, plant height and ear height, and recorded flowering time for a subset (12/32) of mutant alleles during one field season. Traits were measured in two environments that represented distinct planting dates in the same growing season. While there were some examples in which specific mutant alleles exhibited a significant difference in plant or ear height relative to the W22 control in one environment, there were no mutants with a significant difference in both environments (Data File S2). This suggests that these mutants have a limited effect on plant or ear height. The analysis of relative flowering time in both environments revealed that one of the mutant alleles, *c3h42-m1*, shows a delay in tassel shedding by 7 days and ear silking by 3 days (Data File S2). The other mutants did not exhibit significant changes in flowering time in these environments. Overall, there may be subtle phenotypic effects of these mutants but there are very few major qualitative or quantitative morphological differences that are obvious.

As some of the TFs were selected from the initial Y1H screen as putative phenolic biosynthesis regulators, we performed targeted liquid chromatography-mass spectroscopy (LC-MS) for 24 phenolic compounds on all 32 mutant alleles. Although we back-crossed these mutant alleles to the W22 *r-g* inbred, the UniformMu W22 lines include introgression of several loci providing anthocyanin expression. To control for potential segregating variation from these introgression regions we compared the mutant metabolite profiles to controls from both the W22 *r-g* colorless and UniformMu color-converted W22 lines (McCarty *et al*. 2005).

In total, we found that at least 1 mutant allele for 7 (*BAF60.21*, *BZIP76*, *E2F19*, *HSF13*, *HSF24*, *JMJ13* and *MYB40*) of the 13 TFs predicted to regulate phenolic genes by Y1H exhibited statistically significant variation in at least 1 of the 24 phenolic compounds analyzed compared to both W22 controls (Figure 4, Data File S3). There were five phenolic compounds (caffeic acid, maysin, rhamnosylisoorientin, syringic acid, and vanillic acid) that exhibit significant down-regulation in the mutant genotype compared to W22. For two Y1H predicted TFs with two independent mutant alleles, *E2F19* and *MYB40*, the same phenolic compound is significantly down-regulated in both alleles per TF: syringic acid in *EF19* alleles, *e2f19-m1* and *e2f19-m2*, and vanillic acid in *MYB40* mutant alleles, *myb40-m1* and *myb40-m2* (Figure 4, Data File S3). In the initial Y1H screen, *E2F19* showed Y1H interactions with three lignin biosynthesis genes (*COMT1*, *CCR1* and *HCT11*) upstream of syringic acid production and MYB40 exhibits Y1H interactions with six lignin biosynthesis genes (ALDH1, *CCR-like7*, *CCR1*, *COMT1*, *FAH1*, and *HCT6*) upstream of vanillic acid production (Figure S5) (Yang *et al*. 2017). In addition, *OMT1*, another lignin biosynthesis gene, was significantly down-regulated in both E2F19 mutant alleles, *e2f19-m1* and *e2f19-m2*, but was not a Y1H target (Figure S5).

**Figure 4.**
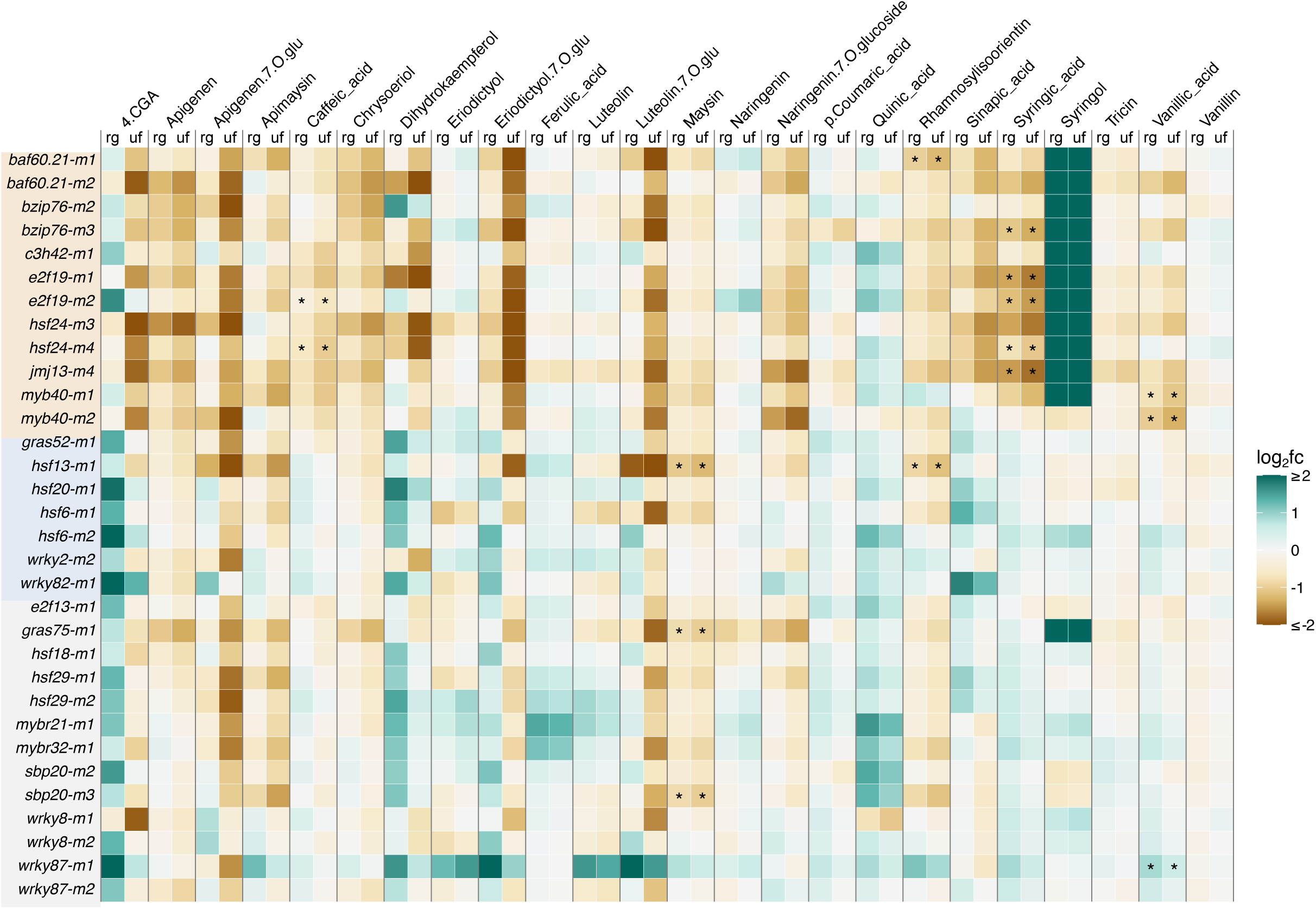
Metabolic profiles of 24 phenolic compounds in TF mutant genotypes. Targeted LC-MS was used to quantify arbitrary units of area (AUA) for 24 phenolic compounds in 32 TF mutant genotypes and 2 W22 control lines: W22 r-g colorless and UniformMu color-converted. Mutant phenolic profiles are shown as the fold change of log2 transformed (log2fc) mutant AUA compared to the AUA of each W22 control, W22 r-g or UniformMu W22 (depicted in figure columns as “rg” for W22 r-g and “uf” for UniformMu W22). Phenolic compounds that are statistically significant (unpaired t-test, FDR adjusted p < 0.05) in a mutant genotype compared to both W22 controls, “rg” and “uf”, are labeled with an asterisk (*). Alleles are grouped by PDI predictions: Y1H (tan), GCN/Y1H (blue), GCN (gray), in the same order as Figure 1C.

### TF mutant allele DE genes reveal enriched GO terms

A potential broader functional role of the TFs was investigated by monitoring enrichment for gene-ontology (GO) terms in each set of DEGs (Wimalanathan *et al*. 2018). There were a relatively large number of GO terms with statistical significance for each mutant allele (Data File S4). To compare potential functional enrichments for the different mutants, we identified the non-redundant set of 50 GO terms with the most significant enrichment for up- or down-regulated DEGs among all 32 mutant alleles (Figure 5, Figure S3). There are several examples in which the two independent mutant alleles for a TF or mutant alleles for TFs in the same family exhibited consistent enrichments. For example, the two mutant alleles of *E2F19* exhibit enrichment for down-regulation of methyl esterase activity and UDP-glycosyltransferase (UGT) activity (Figure 5). The two mutant alleles of *WRKY87* and one allele for *WRKY82* were enriched for down-regulation of mini-chromosome maintenance (MCM) and THO complex genes (Figure 5). There were several GO terms that exhibited significant enrichment in at least eight of the mutants, including UDP-glycosyltransferase (UGT), THO complex, MCM complex, DNA unwinding, and cellular glucan metabolism. Although some mutants exhibited functional enrichment of DEGs, the overall minimal enrichment signal for a particular molecular function or biological process within mutant DEG sets is consistent with plants that display no major phenotypic differences under standard growth conditions.

**Figure 5.**
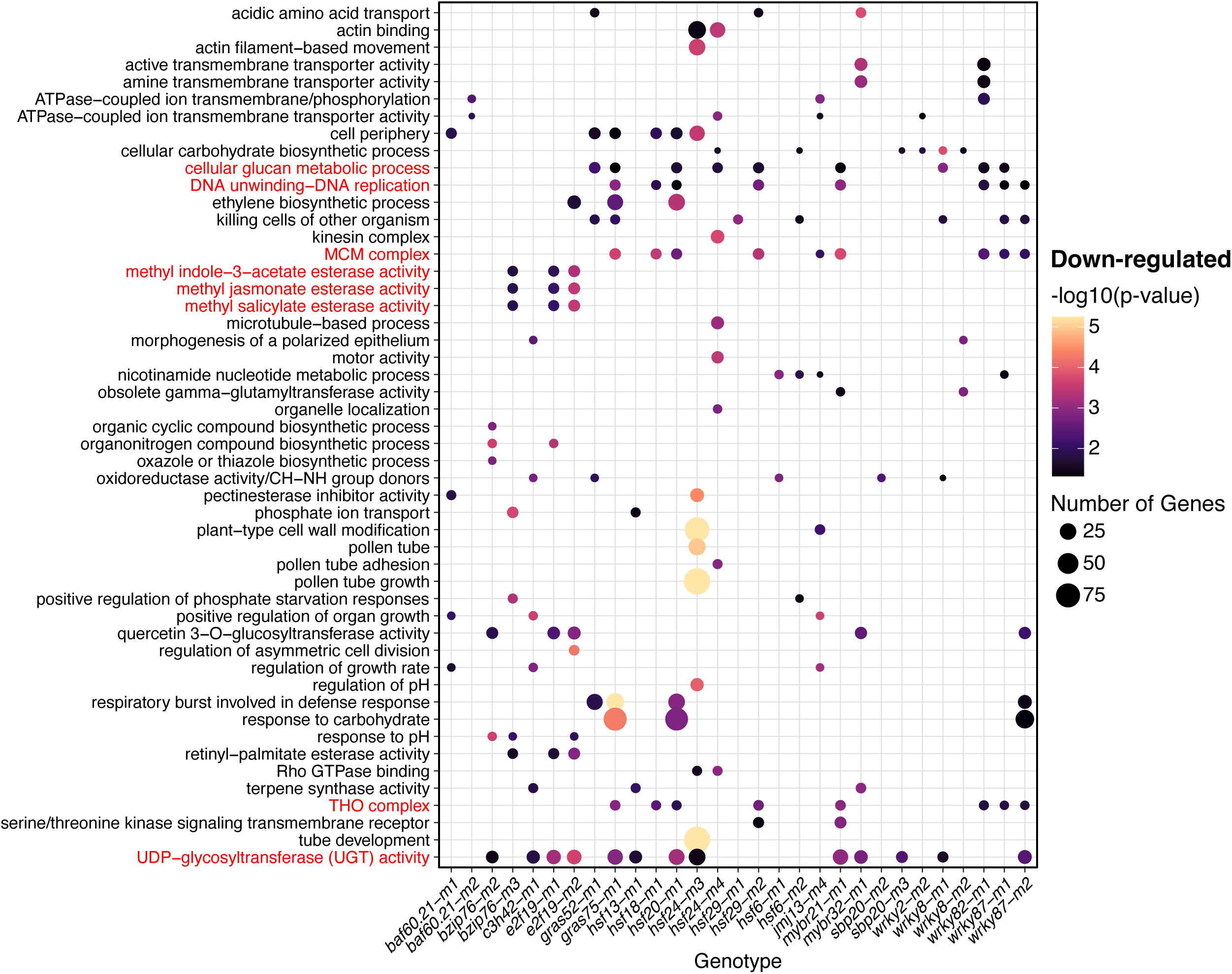
GO-based analysis of TF mutant down-regulated differential expression. A GO analysis of the down-regulated genes for all 32 mutants was used to identify a set of the top 50 non-redundant enriched terms (lowest p-values). Terms highlighted in red are referred to in the text. The GO enrichment levels were determined by a hypergeometric test, where the GO term observed/expected number for each gene set was tested. Circles are used to indicate each significant enrichment of GO terms for 29 of the 32 mutants (*e2f13-m1, myb40-m1*, and *myb40-m2* have no significant enrichments in the top 50 GO terms for down-regulated DEGs).

### A small number of Y1H predicted targets exhibit differential expression

A subset of the TFs (13/22) with mutant alleles have targets predicted from the Y1H screen. These include five TFs with three or less predicted targets and eight TFs with at least four predicted targets (Figure S4, Data File S5). There are some predicted Y1H targets that are not expressed in the profiled tissue or that lack a W22 gene annotation (Y1H targets were predicted for B73v4 based gene annotations) and these could not be tested for altered expression in the mutants (Figure 6, Figure S5, Data File S5). Most of the predicted targets that were expressed did not show evidence for differential expression in the mutants (Figure 6, Figure S5). There were only four significantly down-regulated and seven significantly up-regulated Y1H predicted target genes out of more than 100 predicted targets tested for differential expression. Given the relatively small number of targets for each mutant allele, it was difficult to perform a formal significance analysis, but our results provided some initial evidence that removing a single TF has minimal functional consequences for the expression of most Y1H predicted targets. The analysis between multiple alleles for the same TF did reveal one potentially interesting case of confirmed functional effects for Y1H predicted targets. The gene *Bz1* (*Zm00001d045055;* a UGT involved in the glycosylation of anthocyanidins) is a predicted target of *HSF24* and is significantly down-regulated in both *hsf24-m3* and *hsf24-m4* (Figure 6, Figure S6). The *Bz1* gene contains two predicted HSF binding sites located in the Y1H cloned promoter region. The *E2F19* predicted target gene *HCT11* (*Zm00001d020530*) is significantly down-regulated in *e2f19-m1* and is down-regulated in *e2f19-m2* but not significantly DE (Figure S6). The *MYB40* predicted target gene *A1* (*Zm00001d044122*) has five MYB transcription factor binding sites within the Y1H cloned promoter region and is significantly up-regulated in *myb40-m1 but* is not significantly DE in *myb40-m2* (Figure S6).

**Figure 6.**
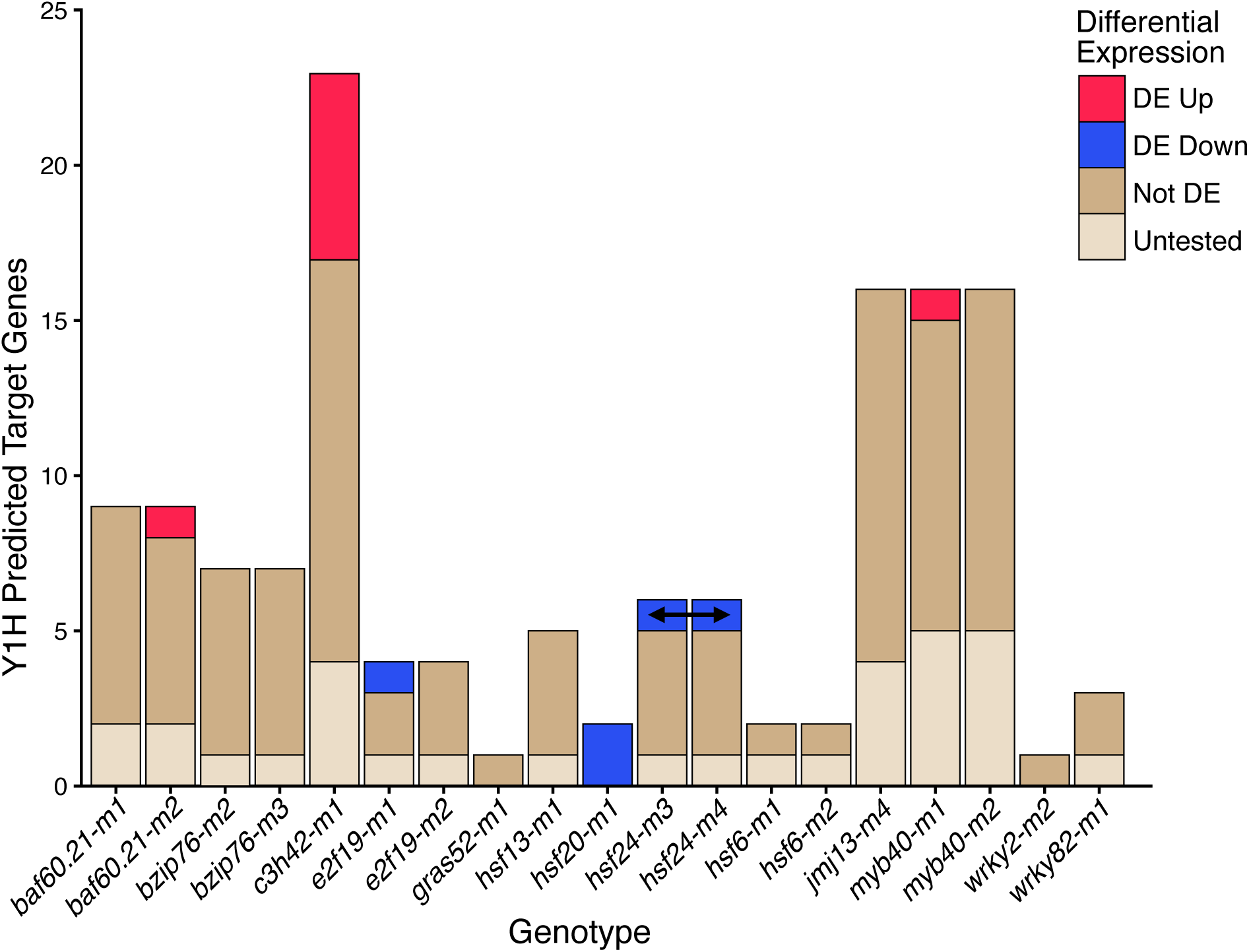
Limited examples of differential gene expression for targets predicted by yeast one-hybrid. The expression of each of the putative targets of each TF that were identified in Y1H analyses (Data File S2) was assessed in the mutants. Each predicted target was classified as DE (Up or Down), not DE or untested. The untested genes include genes that did not have an annotation in the W22 genome and genes that are not expressed in this tissue. The predicted target (*Bz1*, Zm00004b031704) that is significantly down-regulated in both *hsf24* mutant alleles is indicated by an arrow.

### Some TF mutants exhibit enriched differential expression of GCN predicted targets

The GCN predictions provided a much larger set of predicted targets for most of the TFs in this study (Figure S4, Data File S5). Our analysis primarily focused on the full set of predicted targets that are identified in at least one of the GCNs (referred to as n1), but for some analyses we also assessed enrichments for targets that were found in at least three of the GCNs (referred to as n3). For GCN predicted TFs, we observed 13 out of the 29 alleles exhibited significant enrichment of predicted targets in the set of DEGs (Figure 7A). There are three mutant alleles with significant enrichment for only down-regulation, four mutant alleles with significant enrichment for only up-regulation, and six mutant alleles (*hsf18-m1*, *hsf20-m1*, *hsf6-m1*, *mybr32-m*1 and *wrky82-m1*) with significant enrichment for both up- and down-regulated target genes (Figure 7A). In most cases, the level of enrichment for targets is only 1.5 to 2-fold (Figure 7A). While there is a significant enrichment for differential expression of predicted target genes for some of the TFs, the majority (> 80%) of the GCN predicted targets did not exhibit differential expression in the mutant relative to wild type (Figure 7B). A similar set of analyses were performed after restricting the predicted targets to genes identified in at least three of the GCNs, n3 (Figure S7). For three TFs (*HSF18*, *HSF20*, and *WRKY82*), the targets predicted in at least three GCNs, n3, exhibit higher enrichment for differential expression than those predicted in at least one GCN, n1 (Figure S7A, Figure 7A), but most predicted targets did not exhibit altered expression (Figure S7B).

**Figure 7.**
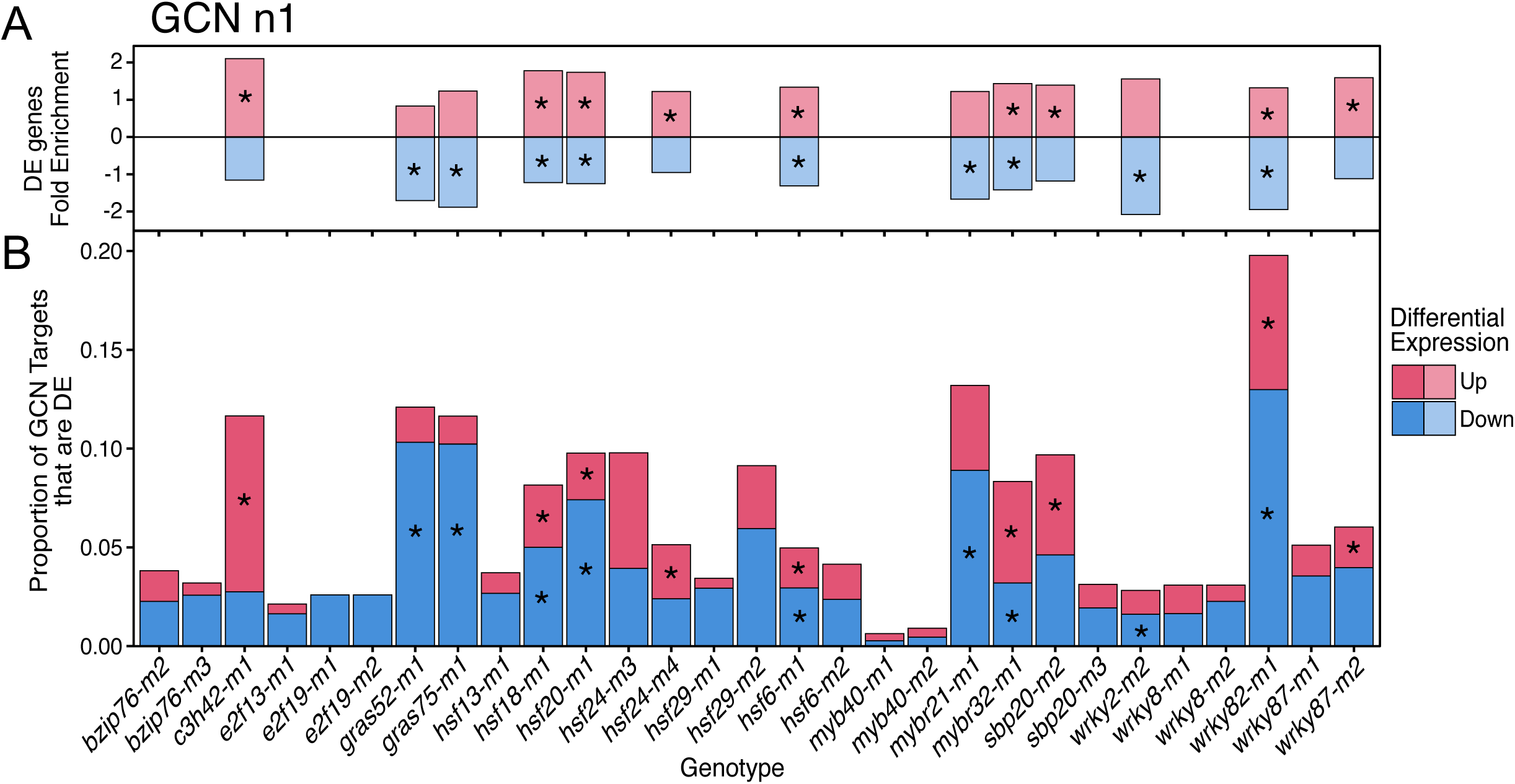
Enrichments of gene co-expression network (GCN) n1 predictions for some TF mutants. The predicted targets for each mutant were identified based on co-expression interactions detected in at least 1 of the 45 GCNs (GCN n1) (Data File S2). **A)** For each mutant allele we calculated the fold-enrichment of DE genes that were predicted targets (the observed number of GCN n1 predicted target genes that were significantly DE divided by the expected number of DE predicted targets) and the enrichments are shown for any mutants that exhibit significant enrichment (*, p < 0.05). **B)** The proportion of GCN predicted targets that are up- (red) or down-(blue) regulated in each mutant is shown. For each allele, significant hypergeometric enrichment for up-regulated (red) and/or down-regulated (blue) DE target genes were marked (*).

## Discussion

Perturbing GRNs is considered a potential mechanism to influence traits in many species. Substantial investments have been made in developing GRNs to predict the functional targets for many TFs and to generate mutant collections for maize genes. In this study, we monitored the consequences of mutant alleles for 22 maize TFs within predicted GRNs. These TFs were selected based on prior Y1H work to identify TFs that bind to promoters of multiple genes in the phenylpropanoid pathway (Yang *et al*. 2017) or based on GCN analyses (Zhou *et al*. 2020a). All the mutant lines that were analyzed are derived from the UniformMu population and are in a W22 genetic background (McCarty *et al*. 2018). Each mutant line was grown in at least two field seasons with replicated plots. No obvious morphological differences were observed in segregating rows comparing mutant and wild type siblings or in plots of homozygous mutants compared to standard W22. The analysis of quantitative height and flowering time traits revealed limited impact for these mutations as well. Overall, the loss-of-function for the single TFs used in this study did not result in major phenotypes in the field environments. In contrast, targeted metabolic profiling of 24 phenolic compounds did reveal that the absence of a single functional TF may significantly change phenolic accumulation. This was most evident for two TFs, *E2F19* and *MYB40*, that were predicted to be regulators of phenolic biosynthesis in the Y1H screen and exhibited significant phenolic compound down-regulation in both independent mutant alleles.

Transcriptome profiling was utilized for three distinct purposes in this study: evaluation of the mutant allele transcripts, GO enrichment analyses of genes with altered expression, and assessment of expression changes for GRN predicted targets. We initially performed transcriptome profiling on a larger set of UniformMu mutant alleles; however, for this study we focused on the subset of alleles that most likely represent loss-of-function mutations. Evaluation of TF gene expression levels in the mutant alleles revealed some cases of reduced expression, but most genes did not exhibit a significant change in transcript abundance. In addition, most mutant alleles had similar patterns of mapped RNA-seq read coverage in regions of the TF gene both 5’ and 3’ of the *Mu* insertion compared to W22. However, the mutants often had reduced coverage in regions flanking the *Mu* insertion site, which is expected if the mutant transcripts include sequences that are novel relative to the W22 reference genome (Ellison *et al*. 2023). To identify potential loss-of-function alleles, we performed *de novo* transcriptome assemblies to determine the mutant transcript structure and sequence. We found that for most of the mutant alleles with a *Mu* insertion in the coding sequence there are often multiple transcripts generated (Ellison *et al*. 2023). Mutant alleles were selected as loss-of-function mutations if there was transcript assembly evidence that the original full-length protein could not be encoded.

Investigating enrichment of GO terms in genes that are DE can be useful in understanding functional consequences of the mutant alleles. Many of the mutants exhibited significant enrichments for some GO terms in the up- or down-regulated DE genes. However, a specific biological function of the TFs could not be determined based on the significantly enriched GO terms.

The evaluation of transcript abundance for genes that are predicted as targets for each of the TFs revealed some examples of altered expression. In about a third of the TFs with GCN predictions there was a significant enrichment of the predicted targets within the DEGs. However, the majority of the DEGs for any specific mutant line are not predicted targets and most of the predicted targets are not DE. We considered three main explanations for this observation. One potential explanation is that the GRN predictions have a high rate of false positives. The types of data used to generate the GRN predictions in this study are both known to have false positives. Y1H assays are conducted in the absence of the normal chromatin environment of the endogenous promoters and can generate false positives. Co-expression analyses are simple guilt-by-association approaches and can suggest functional interactions for sets of genes that are co-expressed in similar patterns, even if these genes have independent regulation. In addition, the GCNs that were used to make predictions were largely based on RNA-seq data from B73 or large panels of maize diversity and these may be less effective at predicting interactions in W22. While these false positives can occur in both types of GRN predictions, Y1H and GCN, we are not confident that this is a primary explanation for the low validation rates we observed. A second explanation is that the TF plays a secondary role in regulation of the predicted target gene resulting in minor changes to gene expression, at least under some conditions. Transcriptional regulation of gene expression can be controlled by large complexes of > 50 TFs with TFs directly or indirectly tethered to the target gene promoter (Nakagawa *et al*. 2018; Göös *et al*. 2022). Some of the TFs within the complex will play a primary role in either bringing the complex to DNA or keeping the complex together, but many of the TFs have secondary roles that may involve fine-tuning of gene expression (Mouchiroud *et al*. 2014; Parab *et al*. 2022). These minor changes in gene expression would have been missed with the criteria used to identify DEGs between the TF loss-of-function alleles and wild type in this study. A third explanation is the potential for functional redundancy in the regulation of these predicted targets. This redundancy could be due to highly similar TFs (either retained duplicated genes from the recent whole-genome duplication event in maize or other members of the same TF family) or other TFs that independently regulate the same target gene. It is worth noting that, in many cases, there are substantial numbers of DEGs in the single TF knockouts, so along with the possibility that there is partially redundant regulation of target genes there must be some non-redundant function of these TFs. The explanation of redundancy could explain why some of the predicted targets are not DE, but we often did not find evidence from Y1H or GCN data that would have implicated other highly similar TFs. This might suggest that the GRN predictions tend to highlight specific potential TF-target interactions when the biological reality might be much more complex. In most cases, we were not able to recover loss-of-function alleles for multiple related TFs, which reduces our ability to perform targeted analyses of redundant regulation. Future studies that utilize genome editing or other approaches to create loss-of-function alleles in multiple TF family members could better explore the potential redundancy of regulation in these pathways.

## Methods

### Isolation of TF mutant alleles from the UniformMu population in maize

Methods for isolating 32 mutant alleles representing 22 TFs are detailed in Ellison et al. 2023 (Ellison *et al*. 2023).

### RNA-seq plant samples, data processing, and transcriptome profiling

Ellison et al. 2023 methods (Ellison *et al*. 2023) detail how: plant material from the 32 TF mutant alleles and W22 r-g control was sampled for RNA-seq, RNA-seq data was processed, transcriptome assembly was performed, and transcriptome profiling was used to identify TF loss-of-function alleles. Our experimental design focused on collecting mutant and wild type transcriptome data from tissues with moderate-high expression (> 6 CPM) of the TF relative to other tissues. In total we surveyed five different tissues: coleoptile tip, imbibed embryo, seedling leaf, tassel, and tassel stem, with variable numbers of mutant alleles assessed (Table S1 and Supplementary Table 2 in Ellison et al. 2023 (Ellison *et al*. 2023)). PCAtools (Blighe and Lun 2023) was used to conduct a principal component analysis of the CPM values of expressed genes and explore sample cluster patterns among and between biological replicates of mutant alleles and wild type W22 control. TF mutant alleles were characterized as potential loss-of-function alleles by determining that variation in the de novo assembled transcript structure and sequence resulted in an allele that could not produce the full-length protein. The 32 TF loss-of-function alleles produced transcripts 5’ of the *Mu* insertion with the initial AUG out-of-frame and/or transcripts 3’ of *Mu* that would only produce a protein less than half the length of the normal protein.

### Identification of TF mutant allele DE genes

Raw read counts of expressed genes (CPM ≥ 1 in at least 1 sample per tissue) from all replicates of each TF mutant allele and W22 control from the same tissue were used to call differentially expressed (DE) genes: false discovery rate (FDR) adjusted p-value < 0.05 and a minimum fold change of 2 (DESeq2 v1.30.1 (Love *et al*. 2014)). To determine if any genes could be DE due to genetic differences between the control W22 r-g colorless and UniformMu color-converted W22 (McCarty *et al*. 2005), DE genes were called between all TF mutant allele and W22 samples from the same tissue. A small number of genes (n = 1773) that are consistently DE in more than half of the mutant alleles from imbibed embryo, tassel stem and seedling leaf tissues (< 50% of mutant reads are from one TF gene) were removed. DE genes for each TF mutant allele were then filtered for expressed genes (CPM ≥ 1 in at least one sample per tissue) and genes with one-to-one gene models (Monnahan *et al*. 2020) between W22 (Springer *et al*. 2018) and B73v4 genomes (Schnable *et al*. 2009; Jiao *et al*. 2017). Filtering for W22 mapped genes with one-to-one B73v4 gene models was necessary for testing enrichment of predicted B73 mapped co-expression-based GRN and Y1H TF targets.

### Morphological field measurements

All mutant alleles were visually assessed for morphological phenotypes across multiple growing seasons in segregating and fixed homozygous rows. Plant height, ear height, and flowering time were measured during one field season for 12/32 mutant alleles in two different fields planted two weeks apart representing distinct environments. Each field consisted of 1 row per mutant allele and 11 rows of W22 control. All trait measurements were only compared within each field to account for any field-related variability. Plant height and ear height were both measured at reproductive maturity on up to 10 non-border plants/row with some variability in the number of plants measured per row due to low stand counts. Plant height was measured as the distance from the soil surface to the base of the flag leaf and ear height was measured as the distance from the soil surface to the highest ear-bearing node. For plant height and ear height (measured in cm), a t-test (unpaired, FDR adjusted p-value < 0.05) was performed on log transformed values between each mutant row and the combined W22 rows within a field to determine if the difference between means was significant (Data File S2).

Flowering time was calculated as the number of days from planting to the tassel shedding or ear silking for at least 50% of the plants in each row (Data File S2). For W22 control, flowering time values were averaged across the 11 rows per field. A significance test was not performed for flowering time as there was only a single value per mutant row in each field replicate. However, if there was greater than +/- 3 days difference in the number of days to tassel shedding or ear silking in a mutant row compared to the combined W22 rows in each independent field environment, there may be a difference in flowering time.

### Phenolic profiling with targeted LC-MS

The 32 mutant alleles and 2 W22 controls, W22 r-g colorless and UniformMu color-converted W22, were grown in 16 h light 28°C, 8 h dark 24°C growth chamber conditions and watered 3 days after sowing and every other day following with 50 ml of water until harvest. Whole shoots were harvested 21 days after sowing (DAS) and 5 shoots were pooled for each of the three biological replicates per mutant or control. At harvest, samples were placed in liquid nitrogen and then were freeze dried prior to lyophilizing. For detailed methods on sample preparation, including tissue extracts, reagent preparation and LC-MS data acquisition and normalization see Materials and Methods sections 3.2, 3.3, 3.4 and 3.6 in Rodriguez et al 2022 (Rodriguez *et al*. 2022). Samples were prepared by combining 50mg of homogenized freeze-dried plant material with an extraction solvent consisting of 80% methanol and 0.1% formic acid. For this experiment, we profiled 31 of the 33 phenolic compounds described in Rodriguez et al 2022, excluding Coniferyl aldehyde and Phenylalanine, using the same available chemical standards and internal standard (Data File S3) (Rodriguez *et al*. 2022). The LC-MS runs were split into two batches due to technical difficulties running all samples in one batch. Due to technical variability between batches, the limits of detection (LOD) for each phenolic compound were measured separately for each batch. The LOD per phenolic compound per batch was set at three times the peak area of the blank (extraction solvent alone) reference value, which was calculated as the mean peak area across blank samples within each batch run (Rodriguez *et al*. 2022). We used the LC-MS relative peak area values per compound instead of absolute values to account for compounds that accumulate at high levels in maize (beyond the upper limit of the external standards). The relative peak area value for each compound per sample was normalized by the weight of the sample to provide arbitrary units of area (AUA). For 20% of the samples in batch 1, two technical replicates per biological replicate were run to determine the reproducibility of the method. The sample biological replicate AUA value was calculated as the mean AUA between the technical replicates. Compounds with more than 20% of the total samples across both batches with AUA values below the respective compound LOD thresholds or more than 10% of sample values missing across both batches were removed from the analysis. In total 3/31 compounds analyzed were removed based on these criteria: Apigenidin, Quercetin, and Shikimic acid.

To determine if the mutant metabolite profile was significantly different from that of W22, an unpaired t-test (FDR adjusted p-value < 0.05) was performed on log2 transformed AUA values of 28 phenolic compounds between each mutant allele and control group: W22 r-g or UniformMu W22, and between control groups (Data File S3). The 4 phenolic compounds (Vitexin, Dihydroquercetin, Isoorientin, Kaempferol) that were significantly different between the control groups, W22 r-g colorless and UniformMu color-converted W22, were removed from the analysis to avoid false positives. In total, we could analyze data for 24 phenolic compounds. We only considered a mutant metabolite profile per compound to be significantly different from the control if there was statistical significance compared to both W22 controls: W22 r-g and UniformMu W22.

### Identifying TF binding sites in Y1H promoter cloned regions

Transcription factor binding sites (TFBSs) were identified in Y1H bait cloned promoter sequences of predicted targets (Yang *et al*. 2017) for two TFs, *MYB40* and *HSF24*. The B73v4 maize genome was scanned for TFBS sequence of five R2R3 MYB TFs (*AtMYB52*, *AtMYB59*, *AtMYB46*, *AtMYB111*, *AtMYB55*) (Franco-Zorrilla *et al*. 2014) and a generic HSF TFBS (5’ *NGAANNTTCN* 3’) (Perisic *et al*. 1989), with N nucleotide weighting set to the GC content of maize genomic DNA to find significant matches (p < 0.01) (FIMO tool in the MEME suite (Grant *et al*. 2011)). These putative genome-wide TFBSs were then subset to Y1H bait promoter cloned regions for predicted targets of MYB40 or HSF24 to identify if these promoter sequences contained the respective TFBSs (BEDTools intersect (Quinlan and Hall 2010)).

### Enrichment for shared DEGs between multiple independent alleles per TF

A significant hypergeometric enrichment of finding more than the expected number of overlapping DEGs between two independent mutant alleles per TF required a representation factor > 1 and p < 0.05 (Lund), where the expected number of genes = (number of genes in allele A × number of genes in allele B) / number of expressed genes. The number of possible shared DEGs between two independent mutant alleles per TF is the minimum number of either up- or down-regulated DEGs between both alleles. The number of shared DEGs was calculated as a proportion out of the possible shared DEGs between two independent alleles per TF (Figure S2).

### Enrichment for TF predicted targets

All statistical analyses for enrichment of DEGs utilized a hypergeometric probability. Testing for over-representation or enrichment of mutant allele DEGs for GO terms (Data File S4) or GCN predicted targets was calculated with R stats v4.0.2 hypergeometric phyper(q = x - 1, m, n = N - m, k, lower.tail = FALSE) function, p < 0.05 for significance ((R Core Team 2020). For GO term enrichment: N = number of expressed genes associated with any GO term, m = number of genes with a specific GO term, k = number of DEGs with any GO term, x = number of DEGs with a specific GO term. For GCN (n1 and n3) predicted target enrichment: N = number of expressed non-redundant predicted target genes in the genome, m = number of expressed predicted targets per TF, k = number of DEGs, x = number of DEGs that are predicted targets. The fold-enrichment of GCN predicted targets that were DE was calculated by [x / (k / N) × m].

### Data and code availability

All supplemental data and scripts necessary for data analysis are available in the GitHub repository at https://github.com/erikamag/GRN_manuscript. The RNA-seq data generated for this study is available at NCBI BioProject: PRJNA936808, titled “Mutator transposon insertions in maize often provide a novel promoter.” Supplemental Table S2 includes meta data for the RNA-seq data used in this study.

## Acknowledgements

We thank the Minnesota Supercomputing Institute at the University of Minnesota (http://www.msi.umn.edu) for providing resources that contributed to the research results reported within this article. We thank Chase Dickson, Hayden Hamsher, Noelle Lynch, Nicole Melamed and Thomas Lidahl for helping with data generation.

---

**Figure S1. Principal component analysis (PCA) of RNA-seq data.**

**Figure S2. Proportion of shared out of possible down- or up-regulated differentially expressed (DE) genes for TFs with multiple independent mutant alleles.**

**Figure S2. Proportion of shared out of possible down- or up-regulated differentially expressed (DE) genes for TFs with multiple independent mutant alleles.**

**Figure S3. GO-based analysis of TF mutant up-regulated differential expression.**

**Figure S4. The number of predicted targets for each TF gene with mutant alleles.**

**Figure S5. Expression changes for phenylpropanoid pathway genes in the TF mutants.**

**Figure S6. Expression differences of three Y1H predicted target genes in TF regulator mutant and wild type genotypes.**

**Figure S7. Enrichments of gene co-expression network (GCN) n3 predictions for some TF mutants.**

**Table S1. 32 Maize TF mutant alleles isolated from the UniformMu population to test GRN predictions.**

**Table S2. TF mutant RNA-seq meta data submitted to NCBI BioProject: PRJNA936808. Data File S1. Differentially expressed genes for each TF mutant allele.**

**Data File S2. Morphological traits measured for a subset of TF mutant alleles.**

**Data File S3. Morphological traits were measured for 12 mutant alleles in two different fields during one field season.**

**Data File S4. Significant hypergeometric enrichment of GO terms associated with TF mutant allele differentially expressed genes.**

**Data File S5. Predicted targets from GCN and Y1H methods for all transcription factors in this study.**

## References

Abnave A., J. John, E. Grotewold, A. I. Doseff, and J. Gray, 2024 Upper level and cross hierarchical regulation of predominantly expressed phenolic genes in maize. Current Plant Biology 39: 100364.

Arda H. E., and A. J. M. Walhout, 2010 Gene-centered regulatory networks. Brief. Funct. Genomics 9: 4–12.

Badia-I-Mompel P., L. Wessels, S. Müller-Dott, R. Trimbour, R. O. Ramirez Flores, et al., 2023 Gene regulatory network inference in the era of single-cell multi-omics. Nat. Rev. Genet. 24: 739–754.

Blighe K., and A. Lun, 2023 PCAtools: everything Principal Component Analysis.

Bolduc N., A. Yilmaz, M. K. Mejia-Guerra, K. Morohashi, D. O’Connor, et al., 2012 Unraveling the KNOTTED1 regulatory network in maize meristems. Genes Dev. 26: 1685–1690.

Brady S. M., L. Zhang, M. Megraw, N. J. Martinez, E. Jiang, et al., 2011 A stele-enriched gene regulatory network in the Arabidopsis root. Mol. Syst. Biol. 7: 459.

Brkljacic J., and E. Grotewold, 2017 Combinatorial control of plant gene expression. Biochim. Biophys. Acta 1860: 31–40.

Burdo B., J. Gray, M. P. Goetting-Minesky, B. Wittler, M. Hunt, et al., 2014 The Maize TFome--development of a transcription factor open reading frame collection for functional genomics. Plant J. 80: 356–366.

Dietrich C. R., F. Cui, M. L. Packila, J. Li, D. A. Ashlock, et al., 2002 Maize Mu transposons are targeted to the 5’ untranslated region of the gl8 gene and sequences flanking Mu target-site duplications exhibit nonrandom nucleotide composition throughout the genome. Genetics 160: 697–716.

Eisen M. B., P. T. Spellman, P. O. Brown, and D. Botstein, 1998 Cluster analysis and display of genome-wide expression patterns. Proc. Natl. Acad. Sci. U. S. A. 95: 14863–14868.

Ellison E. L., P. Zhou, P. Hermanson, Y.-H. Chu, A. Read, et al., 2023 Mutator transposon insertions within maize genes often provide a novel outward reading promoter. Genetics 225: iyad171.

Eveland A. L., A. Goldshmidt, M. Pautler, K. Morohashi, C. Liseron-Monfils, et al., 2014 Regulatory modules controlling maize inflorescence architecture. Genome Res. 24: 431– 443.

Farré G., D. Blancquaert, T. Capell, D. Van Der Straeten, P. Christou, et al., 2014 Engineering complex metabolic pathways in plants. Annu. Rev. Plant Biol. 65: 187–223.

Franco-Zorrilla J. M., I. López-Vidriero, J. L. Carrasco, M. Godoy, P. Vera, et al., 2014 DNA-binding specificities of plant transcription factors and their potential to define target genes. Proc. Natl. Acad. Sci. U. S. A. 111: 2367–2372.

Gaut B. S., M. Le Thierry d’Ennequin, A. S. Peek, and M. C. Sawkins, 2000 Maize as a model for the evolution of plant nuclear genomes. Proc. Natl. Acad. Sci. U. S. A. 97: 7008–7015.

Gomez-Cano L., F. Gomez-Cano, F. M. Dillon, R. Alers-Velazquez, A. I. Doseff, et al., 2020 Discovery of modules involved in the biosynthesis and regulation of maize phenolic compounds. Plant Sci. 291: 110364.

Göös H., M. Kinnunen, K. Salokas, Z. Tan, X. Liu, et al., 2022 Human transcription factor protein interaction networks. Nat. Commun. 13: 766.

Grant C. E., T. L. Bailey, and W. S. Noble, 2011 FIMO: scanning for occurrences of a given motif. Bioinformatics 27: 1017–1018.

Hartwig T., M. Banf, G. P. Prietsch, J.-Y. Zhu, I. Mora-Ramírez, et al., 2023 Hybrid allele-specific ChIP-seq analysis identifies variation in brassinosteroid-responsive transcription factor binding linked to traits in maize. Genome Biol. 24: 108.

Huang J., S. Vendramin, L. Shi, and K. M. McGinnis, 2017 Construction and Optimization of a Large Gene Coexpression Network in Maize Using RNA-Seq Data. Plant Physiol. 175: 568–583.

Ikeuchi M., M. Shibata, B. Rymen, A. Iwase, A.-M. Bågman, et al., 2018 A Gene Regulatory Network for Cellular Reprogramming in Plant Regeneration. Plant Cell Physiol. 59: 765– 777.

Jiao Y., P. Peluso, J. Shi, T. Liang, M. C. Stitzer, et al., 2017 Improved maize reference genome with single-molecule technologies. Nature 546: 524–527.

Johnson D. S., A. Mortazavi, R. M. Myers, and B. Wold, 2007 Genome-wide mapping of in vivo protein-DNA interactions. Science 316: 1497–1502.

Kidwell M. G., and D. Lisch, 1997 Transposable elements as sources of variation in animals and plants. Proc. Natl. Acad. Sci. U. S. A. 94: 7704–7711.

Kuo M. H., and C. D. Allis, 1999 In vivo cross-linking and immunoprecipitation for studying dynamic Protein:DNA associations in a chromatin environment. Methods 19: 425–433.

Lambert S. A., A. Jolma, L. F. Campitelli, P. K. Das, Y. Yin, et al., 2018 The Human Transcription Factors. Cell 172: 650–665.

Li C., Z. Qiao, W. Qi, Q. Wang, Y. Yuan, et al., 2015 Genome-wide characterization of cis-acting DNA targets reveals the transcriptional regulatory framework of opaque2 in maize. Plant Cell 27: 532–545.

Liu S., C. R. Dietrich, and P. S. Schnable, 2009 DLA-based strategies for cloning insertion mutants: cloning the gl4 locus of maize using Mu transposon tagged alleles. Genetics 183: 1215–1225.

Love M. I., W. Huber, and S. Anders, 2014 Moderated estimation of fold change and dispersion for RNA-seq data with DESeq2. Genome Biol. 15: 550.

Lu X., J. Liu, W. Ren, Q. Yang, Z. Chai, et al., 2018 Gene-Indexed Mutations in Maize. Mol. Plant 11: 496–504.

Lund J., Calculation of the representation factor and the associated probability

Macneil L. T., and A. J. M. Walhout, 2011 Gene regulatory networks and the role of robustness and stochasticity in the control of gene expression. Genome Res. 21: 645–657.

McCarty D. R., A. Mark Settles, M. Suzuki, B. C. Tan, S. Latshaw, et al., 2005 Steady-state transposon mutagenesis in inbred maize: Maize steady-state transposon mutagenesis. Plant J. 44: 52–61.

McCarty D. R., P. Liu, and K. E. Koch, 2018 The UniformMu Resource: Construction, Applications, and Opportunities, pp. 131–142 in The Maize Genome, edited by Bennetzen J., Flint-Garcia S., Hirsch C., Tuberosa R. Springer International Publishing, Cham.

Mejia-Guerra M. K., M. Pomeranz, K. Morohashi, and E. Grotewold, 2012 From plant gene regulatory grids to network dynamics. Biochim. Biophys. Acta 1819: 454–465.

Monnahan P. J., J.-M. Michno, C. O’Connor, A. B. Brohammer, N. M. Springer, et al., 2020 Using multiple reference genomes to identify and resolve annotation inconsistencies. BMC Genomics 21: 281.

Morohashi K., M. I. Casas, M. L. Falcone Ferreyra, L. Falcone Ferreyra, M. K. Mejía-Guerra, et al., 2012 A genome-wide regulatory framework identifies maize pericarp color1 controlled genes. Plant Cell 24: 2745–2764.

Mouchiroud L., L. J. Eichner, R. J. Shaw, and J. Auwerx, 2014 Transcriptional coregulators: fine-tuning metabolism. Cell Metab. 20: 26–40.

Nakagawa T., M. Yoneda, M. Higashi, Y. Ohkuma, and T. Ito, 2018 Enhancer function regulated by combinations of transcription factors and cofactors. Genes Cells 23: 808–821.

Parab L., S. Pal, and R. Dhar, 2022 Transcription factor binding process is the primary driver of noise in gene expression. PLoS Genet. 18: e1010535.

Pautler M., A. L. Eveland, T. LaRue, F. Yang, R. Weeks, et al., 2015 FASCIATED EAR4 encodes a bZIP transcription factor that regulates shoot meristem size in maize. Plant Cell 27: 104–120.

Perisic O., H. Xiao, and J. T. Lis, 1989 Stable binding of Drosophila heat shock factor to head-to-head and tail-to-tail repeats of a conserved 5 bp recognition unit. Cell 59: 797–806.

Quinlan A. R., and I. M. Hall, 2010 BEDTools: a flexible suite of utilities for comparing genomic features. Bioinformatics 26: 841–842.

R Core Team, 2020 R: A Language and Environment for Statistical Computing. R Foundation for Statistical Computing, Vienna, Austria.

Riechmann J. L., 2002 Transcriptional regulation: a genomic overview. Arabidopsis Book 1: e0085.

Rodriguez J., L. Gomez-Cano, E. Grotewold, and N. de Leon, 2022 Normalizing and Correcting Variable and Complex LC-MS Metabolomic Data with the R Package pseudoDrift. Metabolites 12. 10.3390/metabo12050435

Scherens B., and A. Goffeau, 2004 The uses of genome-wide yeast mutant collections. Genome Biol. 5: 229.

Schnable P. S., D. Ware, R. S. Fulton, J. C. Stein, F. Wei, et al., 2009 The B73 maize genome: complexity, diversity, and dynamics. Science 326: 1112–1115.

Sperling S., 2007 Transcriptional regulation at a glance. BMC Bioinformatics 8 Suppl 6: S2.

Springer N. M., S. N. Anderson, C. M. Andorf, K. R. Ahern, F. Bai, et al., 2018 The maize W22 genome provides a foundation for functional genomics and transposon biology. Nat. Genet. 10.1038/s41588-018-0158-0

Springer N., N. de León, and E. Grotewold, 2019 Challenges of Translating Gene Regulatory Information into Agronomic Improvements. Trends Plant Sci. 10.1016/j.tplants.2019.07.004

Taylor-Teeples M., L. Lin, M. de Lucas, G. Turco, T. W. Toal, et al., 2015 An Arabidopsis gene regulatory network for secondary cell wall synthesis. Nature 517: 571–575.

Thompson D., A. Regev, and S. Roy, 2015 Comparative analysis of gene regulatory networks: from network reconstruction to evolution. Annu. Rev. Cell Dev. Biol. 31: 399–428.

Tu X., M. K. Mejía-Guerra, J. A. Valdes Franco, D. Tzeng, P.-Y. Chu, et al., 2020 Reconstructing the maize leaf regulatory network using ChIP-seq data of 104 transcription factors. Nat. Commun. 11: 5089.

Walhout A. J. M., 2011 What does biologically meaningful mean? A perspective on gene regulatory network validation. Genome Biol. 12: 109.

Wimalanathan K., I. Friedberg, C. M. Andorf, and C. J. Lawrence-Dill, 2018 Maize GO Annotation-Methods, Evaluation, and Review (maize-GAMER). Plant Direct 2: e00052.

Yang F., W. Z. Ouma, W. Li, A. I. Doseff, and E. Grotewold, 2016 Establishing the Architecture of Plant Gene Regulatory Networks. Methods Enzymol. 576: 251–304.

Yang F., W. Li, N. Jiang, H. Yu, K. Morohashi, et al., 2017 A Maize Gene Regulatory Network for Phenolic Metabolism. Mol. Plant 10: 498–515.

Zhou P., Z. Li, E. Magnusson, F. Gomez Cano, P. A. Crisp, et al., 2020a Meta gene regulatory networks in maize highlight functionally relevant regulatory interactions. Plant Cell 32: 1377–1396.

Zhou P., Z. Li, E. Magnusson, F. Gomez Cano, P. A. Crisp, et al., 2020b Meta Gene Regulatory Networks in Maize Highlight Functionally Relevant Regulatory Interactions. Plant Cell 32: 1377–1396.

